# Batrachotoxin Sensitive Sodium Channels in Toxic Birds Challenge “Target Mutation” Strategy of Toxin Autoresistance

**DOI:** 10.64898/2026.01.29.702343

**Authors:** Evelina Gromilina, Zhiying Jia, Mridhula J Thyagarajan, Hayeon Yang, Dowson Yang, Fayal Abderemane-Ali

**Affiliations:** Department of Physiology, David Geffen School of Medicine, University of California, Los Angeles, Los Angeles, CA 90095, USA; Undergraduate Neuroscience Interdepartmental Program, University of California, Los Angeles, Los Angeles, CA 90095, USA; Department of Chemistry and Biochemistry, University of California, Los Angeles, Los Angeles, CA 90095, USA

**Keywords:** Voltage-gated sodium channel, Batrachotoxin, Toxin resistance, Toxic birds

## Abstract

Voltage-gated sodium channels (Na_V_s) are essential for muscle and nerve activity, and are therefore prime targets of natural toxins. Batrachotoxin (BTX) is uniquely potent among Na_V_-directed alkaloids. Southern and Central American *Phyllobates* poison dart frogs and multiple species of New Guinean toxic birds accumulate dietary BTX without self-poisoning. Two broad autoresistance models have been proposed: 1) “target mutation,” in which Na_V_ substitutions reduce BTX action, and 2) “toxin sequestration,” in which high-affinity binding proteins or compartmentalization limit toxin access to channels. A recent report identified two Na_V_1.4 substitutions, D1050N and S1568P, under positive selection in toxic birds, including two newly discovered BTX-bearing birds, *Pachycephala schlegelii* and *Aleadryas rufinucha*, and suggested a “target mutation” strategy for these passerines. Here, we test that hypothesis using structure-guided mapping and electrophysiology. The substitutions map to solvent-exposed positions tens of angstroms away from inner-cavity BTX sites, inconsistent with direct effects on BTX binding. Heterologously expressed mutant channels exhibit wild-type activation and steady-state inactivation, lacking the biophysical “costs” typical of pore-lining resistance mutations. Importantly, all constructs remain fully BTX-sensitive, showing canonical shifts in activation and persistent currents upon toxin exposure. These data argue that Na_V_1.4 substitutions do not confer BTX autoresistance in toxic birds. This finding, together with prior work in BTX-carrying frogs and birds, is consistent with a generalized sequestration model. Identifying the responsible BTX-binding factors could enable antidote design and broaden strategies for neutralizing Na_V_-targeting toxins.

## Introduction

Voltage-gated sodium channels (Na_V_s) initiate and propagate action potentials and thus underlie neuronal signaling and skeletal/cardiac excitability^1,2^. This Na_V_ central physiological role makes these channels prime targets for natural toxins such as alkaloids and animal venoms that bind to Na_V_s and cause channel dysfunction^2,3^. Among Na_V_-directed toxins, the steroidal alkaloid batrachotoxin (BTX) is exceptional for its potency and mechanism. Unlike pore-blocking toxins such as tetrodotoxin (TTX) and saxitoxin (STX), which cap the outer vestibule to prevent Na^+^ flux, BTX enters the inner cavity, diminishes fast inactivation, and stabilizes the pore in the open conformation, all of which produce a sustained Na^+^ influx, excessive depolarization, and death^4–7^. Mutagenesis and biophysical studies mapped BTX-sensing residues to pore-lining S6 helices, especially in DI/DIII/DIV^8–11^, and recent cryo-EM structures revealed inner-cavity ligand footprints consistent with pharmacology^12^. An evolutionary paradox arises because distantly related vertebrate lineages accumulate diet-acquired BTX while avoiding self-intoxication. These include the neotropical South and Central American poison dart frogs (*Phyllobates*)^13^ and multiple species of Papua New Guinean toxic birds (*Pitohui* spp., *Ifrita kowaldi, Pachycephalidae*, *Oriolidae*, *Ifritidae*, and *Oreoicidae*)^14–16^.

Two broad mechanisms have been proposed for such “autoresistance” in BTX-bearing animals. First, a “target mutation” strategy where substitutions in Na_V_s reduce BTX affinity^11,17^. This target-site resistance has been suggested as the main mechanism of Na_V_-targeting toxin autoresistance in other systems, including preys^18–20^ and predators^21–23^. This idea is supported by elegant examples of TTX- and STX-resistant Na_V_ mutations that commonly entail performance trade-offs for channel activity^18–22,24^. For BTX in particular, a DIVS6 N-to-T substitution in *Phyllobates terribillis* Na_V_1.4 reduces BTX modulation when engineered into rat Na_V_1.4, prompting its proposal as an adaptive change in some poison dart frogs^11,17^. However, this variant is either rare or absent in most BTX-bearing frogs^17,25^ and carries substantial biophysical cost when engineered into bird, human, or rat Na_V_1.4^26^. Surprisingly, despite providing resistance in these dysfunctional channels, the frog DIVS6 N-to-T fails to blunt BTX action in poison-frog Na_V_s in vitro^26^. Therefore, many BTX-bearing birds and frogs express BTX-sensitive Na_V_s despite the frogs themselves being resistant to doses of BTX injection as high as 20 times the lethal dose 50% (LD_50_) based on values for mice^26,27^. These observations indicate that a “target mutation” strategy is insufficient, leading to the idea of upstream protection: “toxin sequestration.” This second strategy involves high-affinity toxin-binding proteins (“toxin sponges”), transporters, or compartmentalization to limit free toxin from reaching Na_V_s in excitable tissues^26^. This sequestration strategy has strong precedents in various species where “toxin sponges” have been identified. These include the Saxitoxin-binding protein Saxiphilin found in the American bullfrog *Rana catesbeiana*^28,29^, the Pufferfish Saxitoxin- and Tetrodotoxin Binding Protein (PSTBP) found in *Fugu pardalis*^30,31^, and the alkaloid-binding globulin found in Dendrobatid poison frogs^32^.

A recent study compared Na_V_1.4 sequences across toxic and non-toxic corvoid passerines and reported two Na_V_1.4 substitutions (D1050N and S1568P) as shared substitutions under positive selection in toxic birds, including two newly discovered BTX-carrying birds, *Pachycephala schlegelii* and *Aleadryas rufinucha*, and suggested these substitutions are adaptive changes conferring BTX autoresistance^16^, raising the question of whether these avian species use the target mutation strategy to withstand their own toxin. Here, we test the proposed avian substitutions, using structural modelling and electrophysiology. We found that both substitutions are solvent-exposed and are located tens of angstroms from the inner-cavity BTX sites, making a direct effect on BTX binding unlikely^10–12^. Further, we found that neither single nor double substitution alters Na_V_1.4 activation or inactivation, arguing against the biophysical “costs” typical of pore-lining resistance mutations^8,9,11,26^. Importantly, all constructs remain fully BTX-sensitive. Hence, our data challenge the hypothesis that BTX autoresistance in toxic birds is based on Na_V_ mutations and instead align with a generalized “toxin-sequestration” model for BTX tolerance across birds and frogs^26,33^.

## Results

### Positions of toxic birds Na_V_1.4 mutations suggest no direct role in BTX binding

To evaluate whether Na_V_1.4 D1050N and S1568P point mutations could affect channel function and/or influence BTX binding, we mapped the corresponding amino acid substitutions onto a predicted AlphaFold model of the BTX-carrying *Pitohui uropygialis meridionalis* Na_V_1.4 (*Pum* Na_V_1.4) channel that we previously cloned^26^. *Pum* Na_V_1.4 is used here as a model because it is the only toxic bird Na_V_1.4-encoding gene with a known full-length sequence. The predicted *Pum* Na_V_1.4 model has an overall per-residue confidence score of 73. This score is a per-residue measure of local confidence, determined by a predicted local distance difference test (pLDDT), with scores scaled from 0 to 100, where higher scores indicate greater confidence and more accurate predictions. This confidence score was particularly high, averaging 84, in the channel transmembrane regions for which multiple structures have previously been solved by cryogenic electron microscopy (cryo-EM) (Figure 1A-B). Such a high score suggests a correct backbone prediction for the transmembrane region, including the BTX-binding sites located in the inner pore cavity and previously defined by mutagenesis and high-resolution cryo-EM studies^8–12^. The predicted *Pum* Na_V_1.4 model also displayed a high structure similarity with the BTX-sensitive human Na_V_1.4 (*Hs* Na_V_1.4) cryo-EM structure^35^, particularly at hallmark Na_V_ features such as the selectivity filter, the voltage sensing domains, the pore domain, and the isoleucine-phenylalanine-methionine (IFM) motif responsible for fast inactivation^2^ (Figure 1C). The two structures can be superimposed with an overall Root Mean Square Deviation (RMSD) of 0.956 Å over 1027 Cα atoms, all corresponding to previously solved regions of the channel. Comparison of the *Pum* Na_V_1.4 model with the BTX-bound rat Na_V_1.5 (*Rn* Na_V_1.5) cryo-EM structure^12^ also showed high similarities, particularly in the transmembrane domains and the pore region where BTX binds (Figure S1A). The two structures are highly comparable with an overall RMSD of 0.783 Å over 1114 Cα atoms. Moreover, between *Pum* Na_V_1.4, *Hs* Na_V_1.4, *Rn* Na_V_1.4, and other toxic and non-toxic species, there is a high conservation of the primary structure of the pore lining S6 transmembrane domains, including residues interacting with BTX in the *Rn* Na_V_1.5-BTX structure^12^ (Figure S2). These observations suggest that *Pum* Na_V_1.4 can be used as a reliable model to accurately map BTX binding sites in the context of toxic bird Na_V_1.4.

**Figure 1:**
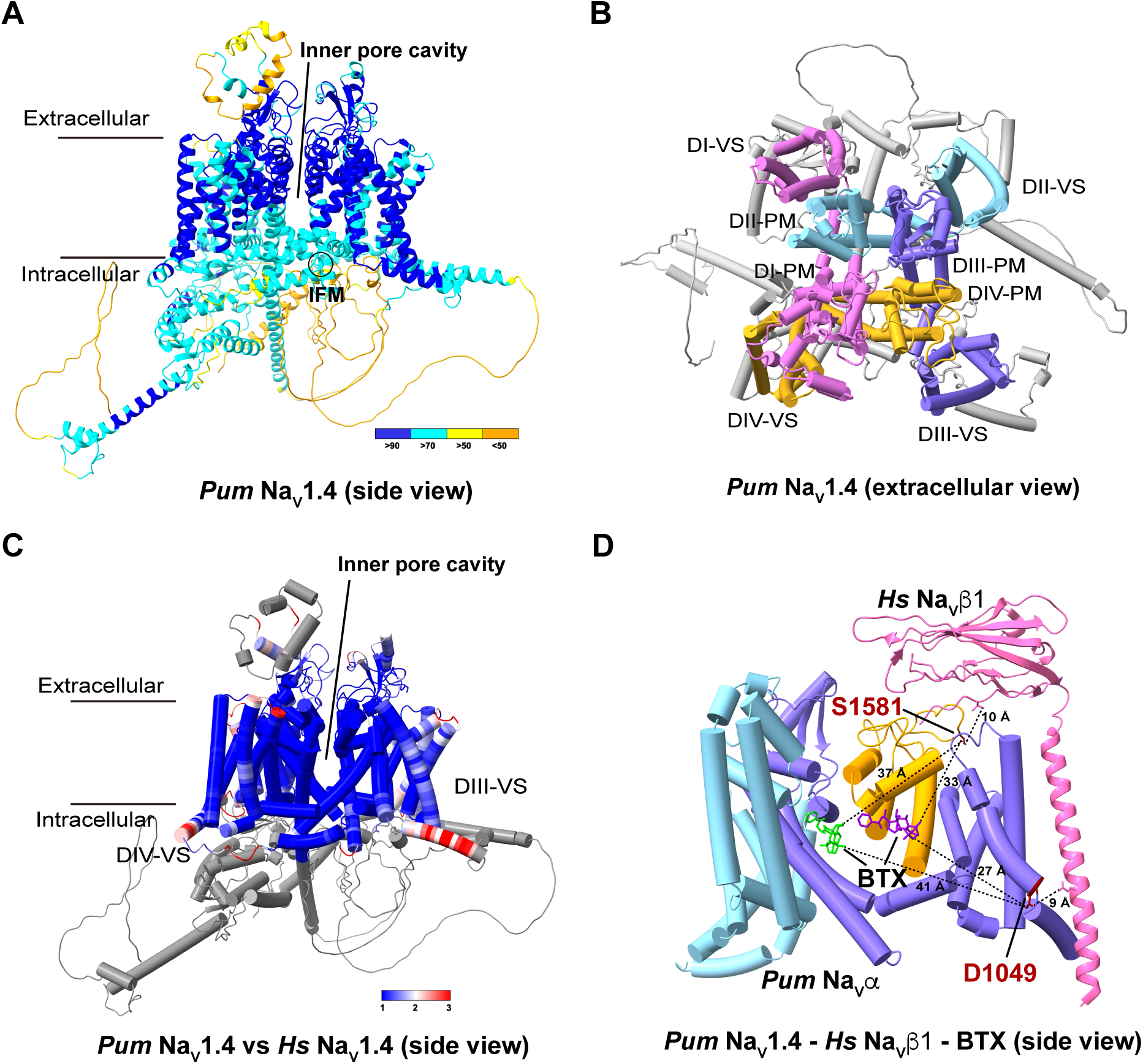
Positions of two Na_V_1.4 mutations in toxic birds are distant from BTX binding sites. **A,** Alphafold predicted structure of *Pum* Na_V_1.4. Side view of overall structure is shown and colored per-atoms with confidence score (pLDDT<50 in orange, 50-70 in yellow, 70-90 in cyan, and >90 in blue). The inner pore cavity and isoleucine-phenylalanine-methionine (IFM) motif are indicated. **B,** Extracellular view of predicted structure of *Pum* Na_V_1.4. DI, DII, DIII, and DIV are colored pink, cyan, purple, and yellow, respectively. Four voltage-sensing (VS) domains and pore modules (PM) are highlighted. **C,** Structural comparison between *Pum* Na_V_1.4 and *Hs* Na_V_1.4. Using *Hs* Na_V_1.4 as a reference, *Pum* Na_V_1.4 is colored by per-residue root mean square deviation (RMSD) with a scale ranging from 1 Å (blue) to 3 Å (red). Regions missing in the experimental *Hs* Na_V_1.4 structure but present in the predicted *Pum* Na_V_1.4 structure are in light gray. The model shows as “Tube Helices”. **D,** Structural model of *Pum* Na_V_1.4-BTX*-Hs* Na_V_β1. *Hs* Na_V_β1 is colored in pink. Residues D1049 and S1581 are colored in red and indicated. BTX molecules inside the channel inner cavity are colored in lime and purple. Residues D1049 and S1581 distances to BTX and *Hs* Na_V_β1 are indicated in dashed lines. For clarity, only parts of Domains II, III, and IV are shown, and Domain I is omitted.

Interestingly, the two avian mutations (D1049N and S1581P; *Pum* Na_V_1.4 numbering) previously proposed to protect *Pachycephala schlegelii, Aleadryas rufinucha* and other toxic birds against BTX are located in regions of the channel that are not involved in BTX binding or modulation. D1049N mutation is located in the intracellular side of the voltage-sensing domain III first segment (VSDIII-S1), while S1581P mutation is on an extracellular loop between the fifth and sixth segment on the pore domain IV (Figure 1D). Notably, none of these substitutions were part of or near the two BTX-binding sites recently determined within the pore inner cavity by cryo-EM^12^. D1049N mutation is located 27 and 41 Å away from these BTX binding sites, while S1581P mutation is 33 and 37 Å away from these BTX binding sites (Figure 1D). Moreover, both mutant positions are solvent-exposed, and their intramolecular interaction networks are also far away from the BTX binding sites and other critical parts of the channel that could affect channel biophysical properties (Figure 1D).

Because Na_V_ beta subunits can form complexes with the alpha subunits and control channel functional expression^39^, with Na_V_β1 and Na_V_β4 known to be expressed in skeletal muscles^40^, we expanded our structural analysis to include these beta subunits and assess whether the two avian mutations (D1049N and S1581P) could reorient and position a Na_V_ beta to block BTX binding.

Given that BTX accesses the Na_V_ inner cavity preferentially in the open state via the intracellular side of the pore and is applied intracellularly in our experiments, we focused on Na_V_β1, the β subunit with a transmembrane helix and intracellular tail that is structurally positioned to influence the intracellular architecture of the α subunit (Figure 1D). By contrast, Na_V_β2 and Na_V_β4 primarily contribute extracellular Ig domains in available α-β structures^41,42^, and Na_V_β3 remains structurally unresolved in complex with α subunits (Figures S1B-D). This structural analysis, using human Na_V_β structures as the only available Na_V_β models, is also supported by a decent conservation between bird and mammalian Na_V_β primary sequence (Figure S3). Our analysis showed that the D1049N and S1581P mutations are located 9 and 10 Å away from Na_V_β1 (Figures 1D and S1D).

These observations suggest that these residues are unlikely to directly perturb BTX binding affinity either by changing the BTX binding pocket or reorienting a Na_V_ beta subunit to block BTX access to the channel pore. This spatial inconsistency raised doubts about the proposed functional significance of these substitutions in autoresistance^16^.

### Mutant channels display wild-type (WT) biophysical properties

Previous studies have shown that mutations of specific residues in the Na_V_1.4 pore domain can reduce or eliminate channel sensitivity to BTX^8–11^. These BTX-resistant mutations include a Na_V_1.4 DIVS6 pore-forming helix N-to-T mutation that was identified in poison-dart frog *Phyllobates terribilis*. When inserted in the Rattus norvegicus Na_V_1.4, this N-to-T mutation renders this channel BTX-insensitive^11^. An assessment of the impact of this mutation on channel biophysical properties revealed that this BTX-resistant mutation incurs a dramatic cost, compromising channel function^26^. Such a cost was observed in engineered BTX-resistant mutants^8,9,26^ and is consistent with the general idea of the trade-offs between toxin-resistant mutations and fitness cost^24^. Therefore, we assessed whether D1049N and S1581P mutations altered channel gating. All Na_V_1.4 α subunits (*Hs* and *Pum* WT and mutants) were co-expressed with human Na_V_β1 for electrophysiology. Wild-type and mutant *Pum* Na_V_1.4 channels bearing each or both of the mutations under positive selection in toxic birds were heterologously expressed in *Xenopus laevis* oocytes. Two-electrode voltage-clamp recordings demonstrated that both WT and mutant channels produced robust inward sodium currents with very similar peak amplitudes and gating properties that are generally affected by BTX-resistant mutations^10,11,26^ or BTX itself^43–45^ (Figure 2A, Table 1). Analysis of current-voltage (I–V) relationships revealed no significant difference in activation threshold, slope factor, or voltage for half-maximal activation (*V*_1/2_-I) between wild-type and mutant channels (Figures 2A-B and Table 1). Similarly, steady-state inactivation curves and inactivation voltage-dependence properties were nearly identical (Figure 2C and Table 2). Because changes in kinetics can occur without appreciable shifts in steady-state voltage dependence, we also measured both inactivation time constant and recovery from inactivation in WT and mutant channels. These biophysical properties are also mostly unchanged between *Pum* Na_V_1.4 WT and mutant channels, besides a slight acceleration of the recovery from inactivation in the double mutant (Figure 2 and Table 3). These data indicate that D1049N and S1581P mutations do not affect the intrinsic gating properties of Na_V_1.4 channels. As BTX-resistant mutations generally come with a fitness cost, these mutations are less likely to affect BTX sensitivity.

**Figure 2:**
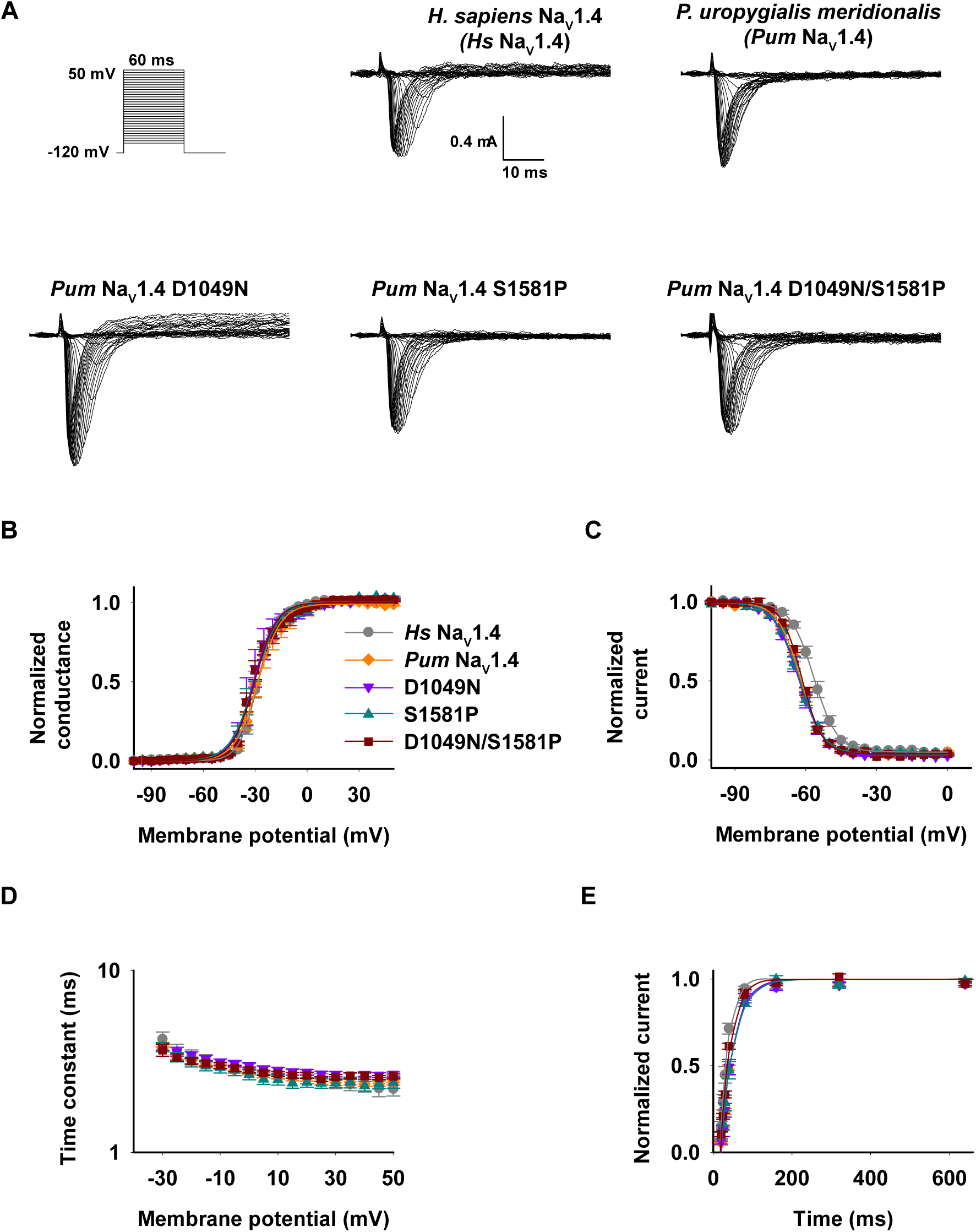
Biophysical characterization of wild-type (WT) and mutant channels. **A,** Exemplar current recordings for *Hs* Na_V_1.4, *Pum* Na_V_1.4, *Pum* Na_V_1.4 D1049N, *Pum* Na_V_1.4 S1581P, and *Pum* Na_V_1.4 D1049N/S1581P expressed in *Xenopus Laevis* oocytes. Currents were evoked with the shown multistep depolarization protocol. **B,** G-V relationships for *Hs* Na_V_1.4 (grey circle), *Pum* Na_V_1.4 (orange diamond), *Pum* Na_V_1.4 D1049N (purple inverted triangle), *Pum* Na_V_1.4 S1581P (green triangle), and *Pum* Na_V_1.4 D1049N/S1581P (dark red square). Solid lines are Boltzmann fits. No significant difference was observed between *Pum* Na_V_1.4 WT and mutants. n = 10–14. **C,** Steady-state inactivation voltage dependence for *Hs* Na_V_1.4 (grey circle), *Pum* Na_V_1.4 (orange diamond), *Pum* Na_V_1.4 D1049N (purple inverted triangle), *Pum* Na_V_1.4 S1581P (green triangle), and *Pum* Na_V_1.4 D1049N/S1581P (dark red square). Solid lines are Boltzmann fits. No significant difference was observed between *Pum* Na_V_1.4 WT and mutants. n = 8–15. **D,** fast inactivation kinetics for *Hs* Na_V_1.4 (grey circle), *Pum* Na_V_1.4 (orange diamond), *Pum* Na_V_1.4 D1049N (purple inverted triangle), *Pum* Na_V_1.4 S1581P (green triangle), and *Pum* Na_V_1.4 D1049N/S1581P (dark red square). No significant difference was observed between *Pum* Na_V_1.4 WT and mutants. n = 10–14. **E,** Recovery from fast inactivation for *Hs* Na_V_1.4 (grey circle), *Pum* Na_V_1.4 (orange diamond), *Pum* Na_V_1.4 D1049N (purple inverted triangle), *Pum* Na_V_1.4 S1581P (green triangle), and *Pum* Na_V_1.4 D1049N/S1581P (dark red square). Solid lines are exponential fits. No significant difference was observed between *Pum* Na_V_1.4 WT and mutants. n = 6–13.

**Table 1:** Activation parameters.

| Channel | Current amplitude (mA) | P-value | $V_{1/2-I}$ (mV) | P-value | $DV_{1/2-BTX}$ (mV) | $k_{act-I}$ (mV) | P-value | $V_{1/2-II}$ (mV) | P-value | $k_{act-II}$ (mV) | P-value | n |
| --- | --- | --- | --- | --- | --- | --- | --- | --- | --- | --- | --- | --- |
| **Hs Nav1.4** | $-1.1 \pm 0.0$ | n.s. | $-28.4 \pm 0.9$ | n.s. | | $4.7 \pm 0.3$ | * | | | | | 14 |
| + BTX | | | $-57.4 \pm 1.5$ | * | $-29.0 \pm 1.8$ | $2.8 \pm 0.4$ | n.s. | $-27.6 \pm 2.1$ | n.s. | $6.3 \pm 0.7$ | n.s. | 15 |
| **Pum Nav1.4** | $-1.0 \pm 0.1$ | n.a. | $-29.0 \pm 1.4$ | n.a. | | $6.5 \pm 0.6$ | n.a. | | | | | 11 |
| + BTX | | | $-61.2 \pm 1.1$ | n.a. | $-32.2 \pm 1.8$ | $2.8 \pm 0.3$ | n.a. | $-31.3 \pm 2.2$ | n.a. | $6.5 \pm 0.8$ | n.a. | 13 |
| **Pum Nav1.4 D1049N** | $-1.1 \pm 0.0$ | n.s. | $-30.8 \pm 1.2$ | n.s. | | $5.8 \pm 0.5$ | n.s. | | | | | 11 |
| + BTX | | | $-65.4 \pm 2.2$ | n.s. | $-34.5 \pm 2.5$ | $3.7 \pm 0.3$ | n.s. | $-35.0 \pm 2.5$ | n.s. | $6.6 \pm 0.9$ | n.s. | 9 |
| **Pum Nav1.4 S1581P** | $-1.1 \pm 0.0$ | n.s. | $-28.8 \pm 0.6$ | n.s. | | $6.5 \pm 0.3$ | n.s. | | | | | 14 |
| + BTX | | | $-64.3 \pm 2.1$ | n.s. | $-35.5 \pm 2.2$ | $4.8 \pm 0.7$ | * | $-31.8 \pm 2.3$ | n.s. | $6.8 \pm 0.8$ | n.s. | 16 |
| **Pum Nav1.4 D1049N/S1581P** | $-1.1 \pm 0.0$ | n.s. | $-30.3 \pm 1.4$ | n.s. | | $5.6 \pm 0.5$ | n.s. | | | | | 10 |
| + BTX | | | $-65.5 \pm 1.0$ | * | $-35.2 \pm 1.7$ | $2.9 \pm 0.5$ | n.s. | $-34.7 \pm 1.8$ | n.s. | $7.7 \pm 0.9$ | n.s. | 13 |

Data are mean ± standard error of the mean.
$V_{1/2-I}$ and $V_{1/2-II}$ are the half-activation potentials. $k_{act-I}$ and $k_{act-II}$ are the slope factors.
Values having "I" only indicate those fit by a single Boltzmann equation.
Values having "I" and "II" indicate cases where the data are fit by a double Boltzmann equation.
P-Values are calculated relative to *Pum Nav1.4* or *Pum Nav1.4* + BTX.
n.a., not applicable.
n.s., not significant; $P > 0.05$ .
"\*" indicates $0.01 < P < 0.05$ .

**Table 2:** Inactivation voltage dependence parameters.

| Channel | $V_{1/2-inact}$ (mV) | P-value | $k_{inact}$ (mV) | P-value | n |
| --- | --- | --- | --- | --- | --- |
| *Hs* Nav1.4 | $-55.9 \pm 0.8$ | *** | $5.3 \pm 0.2$ | n.s. | 15 |
| *Pum* Nav1.4 | $-61.8 \pm 0.8$ | n.a. | $5.2 \pm 0.4$ | n.a. | 9 |
| *Pum* Nav1.4 D1049N | $-62.5 \pm 1.1$ | n.s. | $5.3 \pm 0.3$ | n.s. | 10 |
| *Pum* Nav1.4 S1581P | $-62.3 \pm 0.8$ | n.s. | $5.5 \pm 0.2$ | n.s. | 8 |
| *Pum* Nav1.4 D1049N/S1581P | $-61.6 \pm 0.7$ | n.s. | $5.3 \pm 0.6$ | n.s. | 11 |

Data are mean ± standard error of the mean.
$V_{1/2-inact}$ is the half-inactivation potential.
$k_{inact}$ is the slope factor.
P-Values are calculated relative to *Pum* Nav1.4.
n.a., not applicable.
n.s., not significant; $P > 0.05$ .
“\*\*\*” indicates $P < 0.001$ .

**Table 3:** Inactivation kinetics parameters.

| Channel | Inactivation time constant at -10 mV (ms) | P-value | n | Recovery from inactivation time constant (ms) | P-value | n |
| --- | --- | --- | --- | --- | --- | --- |
| *Hs* Nav1.4 | $3.1 \pm 0.2$ | n.s. | 14 | $33.4 \pm 3.9$ | ** | 13 |
| *Pum* Nav1.4 | $3.0 \pm 0.1$ | n.a. | 11 | $44.5 \pm 1.7$ | n.a. | 7 |
| *Pum* Nav1.4 D1049N | $3.1 \pm 0.1$ | n.s. | 11 | $44.3 \pm 4.3$ | n.s. | 7 |
| *Pum* Nav1.4 S1581P | $2.9 \pm 0.2$ | n.s. | 14 | $45.6 \pm 2.1$ | n.s. | 6 |
| *Pum* Nav1.4 D1049N/S1581P | $3.0 \pm 0.1$ | n.s. | 10 | $37.6 \pm 1.7$ | * | 9 |

Data are mean ± standard error of the mean.
P-Values are calculated relative to *Pum* Nav1.4.
n.a., not applicable.
n.s., not significant; $P > 0.05$ .
“\*” indicates $0.01 < P < 0.05$ .
“\*\*\*” indicates $0.001 < P < 0.01$ .

### Na_V_1.4 D1049N and S1581P Mutant channels retain full sensitivity to BTX

To directly test the hypothesis that the two observed mutations under positive selection in BTX-carrying birds confer BTX resistance, we measured the pharmacological sensitivity of wild-type *Pum* Na_V_1.4, single mutants D1049N and S1581P, and the double mutant D1049N/S1581P, to intracellular application of 50 μM BTX. In these experiments, BTX was applied at a target intracellular concentration of 50 µM to produce partial channel modification, thereby avoiding saturation and enabling detection of potential differences in BTX sensitivity between WT and mutants. Consistent with previous studies^5,7,26^, BTX application on *Hs* Na_V_1.4 and *Pum* Na_V_1.4 resulted in a hyperpolarized shift in the voltage dependency of activation (ΔV_1/2 BTX_ = -29.0 ± 1.8 - 32.2 ± 1.8 mV for *Hs* Na_V_1.4 and *Pum* Na_V_1.4, respectively), and a partial and sustained elimination of fast inactivation (Figure 3 and Table 1). The activation curve under BTX application follows a double-Boltzmann function, with the first and second components arising from BTX-bound and unmodified channels, respectively^10,26^. Notably, comparison of BTX-induced effects on these gating parameters showed no significant differences between wild-type and mutant *Pum* Na_V_1.4 channels (Figure 3 and Table 1). Similar to WT *Pum* Na_V_1.4, BTX shifted activation voltage dependence (ΔV_1/2 BTX_ = -34.5 ± 2.5, -35.5 ± 2.2, and -35.2 ± 1.7 mV for *Pum* Na_V_1.4 D1049N, S1581P, and double mutant D1049N/S1581P, respectively) and partially eliminated steady-state inactivation for all three *Pum* Na_V_1.4 mutants. Importantly, although the fraction of BTX-modified channels varies across oocytes, population-level quantification shows the average BTX-modified fraction is the same for WT and mutants. The extent of inactivation, characterized by the ratio between persistent steady state current during depolarization and peak current, was indistinguishable between *Hs* Na_V_1.4, *Pum* Na_V_1.4 and mutants (Figures 4A-E). Similarly, the BTX-dependent inactivation block was similar, around 30%, between *Hs* Na_V_1.4, *Pum* Na_V_1.4, and mutants, demonstrating equal sensitivity to BTX (Figure 4F). These results strongly suggest that the point mutations found to be under positive selection in the Na_V_1.4 channels of BTX-carrying birds, *Pachycephala schlegelii, Aleadryas rufinucha*, and other toxic birds^16^, do not confer BTX resistance at the level of channel function and therefore, cannot be the BTX autoresistance strategy in these toxic birds.

**Figure 3:**
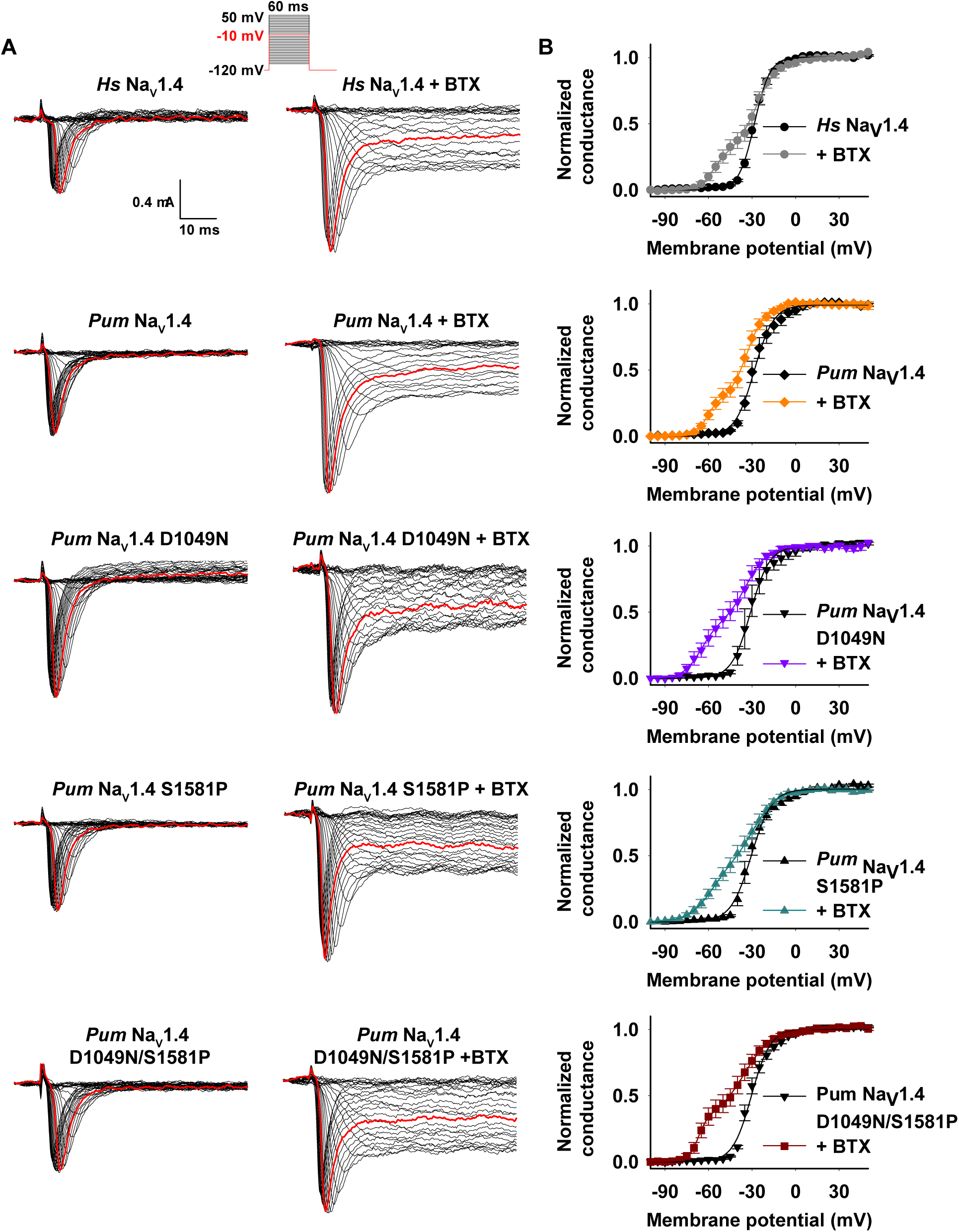
Bird wild-type (WT) and mutant channels are BTX sensitive. **A,** Exemplar current recordings for *Hs* Na_V_1.4, *Pum* Na_V_1.4, *Pum* Na_V_1.4 D1049N, *Pum* Na_V_1.4 S1581P, and *Pum* Na_V_1.4 D1049N/S1581P expressed in *Xenopus Laevis* oocytes before (left) or after (right) intracellular BTX injection (50 nL of 1 mM stock for a target [BTX]i = 50 µM) followed by 1,000 depolarizing pulses at 2 Hz to promote state-dependent access. The trace at 0 mV is highlighted in red in each panel. Currents were evoked with the shown multistep depolarization protocol (inset). **B,** G-V relationships in the presence or absence of BTX for *Hs* Na_V_1.4 (black circle), +BTX (grey circle); *Pum* Na_V_1.4 (black diamond), +BTX (orange diamond); *Pum* Na_V_1.4 D1049N (black inverted triangle), +BTX (purple inverted triangle); *Pum* Na_V_1.4 S1581P (black triangle), +BTX (green triangle); and *Pum* Na_V_1.4 D1049N/S1581P (black square), +BTX (dark red square). Solid lines are Boltzmann fits. n = 9–16.

**Figure 4:**
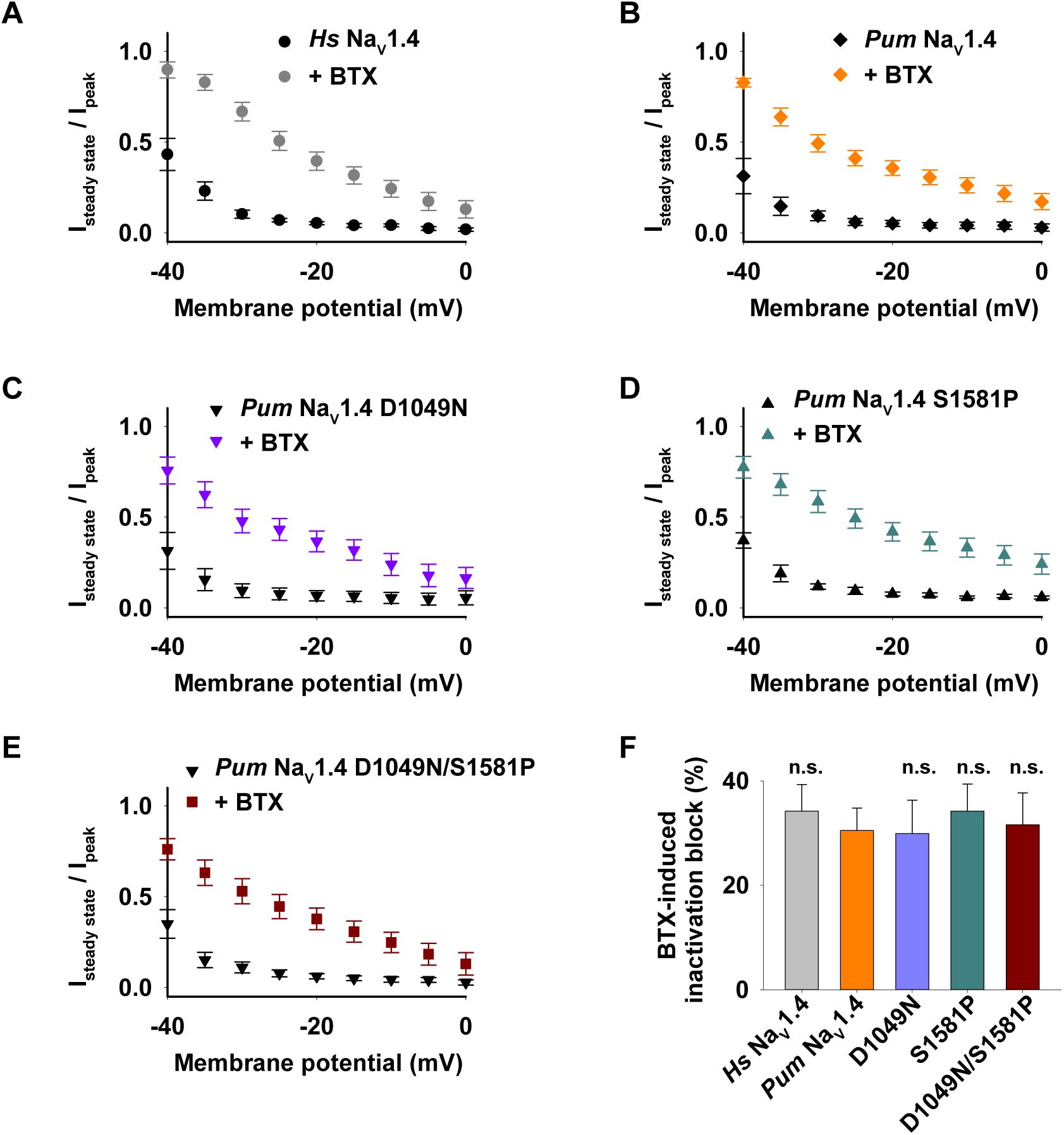
Bird wild-type (WT) and mutant channels have similar BTX-dependent inactivation block. Ratio between persistent steady state current during depolarization and peak current in the presence or absence of BTX for **A,** *Hs* Na_V_1.4 (black circle), +BTX (grey circle); **B,** *Pum* Na_V_1.4 (black diamond), +BTX (orange diamond); **C,** *Pum* Na_V_1.4 D1049N (black inverted triangle), +BTX (purple inverted triangle); **D,** *Pum* Na_V_1.4 S1581P (black triangle), +BTX (green triangle); and **E,** *Pum* Na_V_1.4 D1049N/S1581P (black square), +BTX (dark red square). n = 9–16. **F,** average percentage of BTX-induced inactivation block at -20 mV for *Hs* Na_V_1.4 (grey), *Pum* Na_V_1.4 (orange), *Pum* Na_V_1.4 D1049N (purple), *Pum* Na_V_1.4 S1581P (green), *Pum* Na_V_1.4 D1049N/S1581P (dark red). This percentage of BTX-induced inactivation block at -20 mV depolarization was determined as the difference between I steady state / I peak in the presence and in the absence of BTX. No significant difference was observed between *Pum* Na_V_1.4 WT and mutants. n = 9–16.

## Discussion

Taken together, our structural and functional analyses provide strong evidence that the two Na_V_1.4 point mutations (D1050N and S1568P) identified in *Pachycephala schlegelii, Aleadryas rufinucha*, and other toxic birds do not mediate resistance to BTX. Although these mutations were previously proposed to represent adaptive changes conferring BTX autoresistance in toxic birds^16^, our data contradict this interpretation. Both residues are positioned outside of the BTX-binding cavity, are solvent-exposed, and lie far away from Na_V_ auxiliary β subunits and known BTX-interacting regions identified through mutagenesis and cryo-EM studies^8–12^. In functional assays, neither single nor combined mutations altered channel gating, activation thresholds, or voltage-dependence of activation or inactivation, and most importantly, all variants remained fully sensitive to BTX. Together, these results demonstrate that these Na_V_1.4 point mutations are not responsible for BTX resistance in these avian species.

### Structural mapping does not support direct BTX coordination by the avian substitutions

The central structural observation in this study is that the two Na_V_1.4 substitutions reported in BTX-bearing birds (D1050N and S1568P; corresponding to D1049N and S1581P in *Pum* Na_V_1.4 numbering)^16^ are not positioned within the inner-cavity regions that define BTX receptor sites in mutagenesis and cryo-EM studies^8–12^. Instead, D1049 is located on the cytosolic face of VSDIII-S1 and S1581 resides on an extracellular DIV loop, exposing both residues to either extracellular or intracellular solvent and placing them tens of angstroms from the BTX molecules observed in the Rn Na_V_1.5-BTX structure (Figure 1D)^12^. While mutations outside a ligand pocket can, in principle, modulate toxin action allosterically, we did not directly measure BTX binding affinity. Rather, we infer BTX interaction through its well-established functional signature on Na_V_ gating^7–11,26,46^, which remains indistinguishable between WT and mutant channels in our experiments. Moreover, these D1049 and S1581 residues are located more than 9 Å away from the Na_V_β subunits (Figures 1D and S1). Such positions make it unlikely that these residues participate in BTX binding to, affinity for, or action on Na_V_1.4 channel.

This structural mismatch is important for two reasons. First, it directly challenges the inference made from sequence-based in silico analysis alone that these sites underwent positive selection because of BTX exposure^16^. Although the positive selection is very attractive, it does not necessarily imply functional coupling to a suspected agent or feature. It may reflect a demographic history or selection on another trait in the same protein. Second, our modeling underscores a recurring lesson from Na_V_ toxicology where resistance-conferring mutations that affect toxin affinity are often located on structural “hotspots” formed by the pore-lining helices, selectivity filter, voltage-sensing modules, or inactivation apparatus, all of which play a critical role in channel gating^2^. D1050N and S1568P are not in such hotspots. Therefore, from a structural-function standpoint, these mutations do not make sense as a BTX resistance strategy. Further investigation is needed to find any other functional purpose or evolutionary reason for these mutations.

### Mutations do not incur the biophysical “cost” typical of toxin-resistant Na_V_s

A second line of evidence comes from electrophysiology. Classic BTX-resistance mutations, whether naturally occurring like in *Phyllobates terribilis*^17^ or engineered into mammalian Na_V_1.4, carry an obvious biophysical penalty characterized by either reduced current densities, shifted activation or inactivation voltage-dependence, altered gating kinetics, or destabilized activation or inactivation states^8,9,11,26^. This is consistent with the general idea of the trade-offs between toxin resistance and fitness cost described for multiple toxin-resistant channels and receptors: the more strongly a target is altered to evade toxin binding, the more likely it is to drift away from its optimal activity^24^.

However, in our recordings, *Pum* Na_V_1.4 carrying D1049N, S1581P, or the double substitution behaved like wild-type: voltage dependence of activation and inactivation, slope factors, and inactivation kinetics were all similar to wild-type *Pum* Na_V_1.4. The recovery from inactivation was also unchanged between *Pum* Na_V_1.4 and the single mutants and was only slightly accelerated in the double mutant D1049N, S1581P, a small but statistically significant acceleration, unlikely to affect BTX phenotype. Current amplitudes were also comparably robust in both WT and mutant *Pum* Na_V_1.4. This demonstrates that these two substitutions are biophysically neutral with respect to channel function. On its own, the lack of biophysical change in these mutant channels is not surprising, as channels do accumulate neutral substitutions. But biophysical neutrality is unexpected if a mutation really sits in or transmits to the BTX pockets, because BTX-site residues are parts of the channel inner pore^12^. Therefore, the fact that we see no biophysical changes in the presence of these mutations strengthens the view that these mutations are not acting where BTX acts.

### Full preservation of BTX sensitivity argues against a Na_V_ mutation-based autoresistance

The decisive test was pharmacology. When we challenged WT and mutant channels with intracellular BTX, all constructs showed the canonical signature of BTX sensitivity: a hyperpolarizing shift in activation and a reduced fast inactivation. These BTX effects are similar to those described for BTX-bound Na_V_1.4 and Na_V_1.5^7–11,26,46^. Quantitatively, the BTX-induced shifts in V_0.5_ activation, the fraction of non-inactivating current, and the double-Boltzmann behavior of activation were indistinguishable between WT and any mutant, including the double mutant. If D1049N or S1581P reduced BTX affinity even modestly, we would have expected at least one of the following: (i) a smaller shift in V_0.5_ activation or (ii) a near complete inactivation. None were observed.

This result formally rules out the hypothesis that BTX autoresistance in toxic birds is rooted in these Na_V_ mutations. Unlike the DIV-S6 N-to-T mutation described in *Phyllobates terribilis*^17^, which, when inserted into rat Na_V_1.4, clearly alters BTX action but at a cost^11^, the avian mutations leave BTX action intact. Thus, the toxic birds do not appear to be protecting their Na_V_1.4 channels via target modification. Interestingly, even the DIV-S6 N-to-T mutation described in *Phyllobates terribilis* as a primary BTX-resistance strategy is not enough to protect BTX-carrying frogs. Although this DIVS6 N-to-T mutation alters the BTX responses of bird, human, and rat Na_V_1.4s, it failed to protect poison frog Na_V_s^26^. These observations further challenge the “target mutation” strategy and suggest an alternative toxin autoresistance strategy in BTX-carrying animals.

A caveat deserves to be mentioned in our studies. Our functional tests are performed in heterologous systems (*Xenopus Laevis* oocytes) using human Na_V_β1 rather than avian Na_V_β orthologs. While this approach matches prior functional studies of BTX-bearing bird and frog Na_V_s^26^, it remains possible that species-specific β-subunits, lipid composition, or post-translational modifications modulate Na_V_ pharmacology in vivo.

### A toxin-sequestration model better explains BTX tolerance in birds and frogs

If Na_V_ mutations do not protect the channels from BTX, the defense must occur before BTX reaches Na_V_s in excitable tissues. Therefore, our findings converge with previous studies consistent with “toxin sequestration”, rather than “target mutation”, as the dominant BTX-autoresistance strategy in both poison birds and poison frogs^26^. In this model, the animal carries high systemic or tissue-associated levels of BTX, but the toxin is buffered by BTX-binding proteins (“toxin sponges”), specialized carrier complexes, or subcellular compartmentalization that prevents free BTX from accessing Na_V_s in muscle, heart, brain, or peripheral nerves. This idea is biologically plausible for several reasons: 1) It decouples toxicity from channel constraint. By neutralizing toxin extracellularly or in circulation, the animal does not need to modify a functionally optimized channel like Na_V_1.4, avoiding the fitness costs seen in pore-site mutations^24^. 2) It would allow for the safe transport of the ingested toxin from the digestive system and its concentration in specialized defensive organs such as the skin and feathers^47,48^. 3) It fits with other animals expressing high-affinity “toxin-sponges” such as the Saxitoxin-binding protein Saxiphilin found in the American bullfrog *Rana catesbeiana*^28,29^, the Pufferfish Saxitoxin- and Tetrodotoxin Binding Protein (PSTBP) found in *Fugu pardalis*^30,31^, and the alkaloid-binding globulin found in Dendrobatid poison frogs^32^. A putative toxin-binding protein has also recently been suggested in the pan-Amazonian Royal Ground snake *Erythrolamprus reginae*^49^. All these “toxin-sponges” proteins are known and have been proposed to prevent autointoxication through sequestration^50^. 4) It scales with toxin load as a sequestration strategy can, in principle, match the high BTX content reported in some *Pitohui* and *Phyllobates* species without requiring multiple and coordinated channel mutations.

Our data suggest that *Pachycephala schlegelii* and *Aleadryas rufinucha* are simply additional examples that fall into this sequestration-centric framework. The fact that these birds harbor BTX and do not have BTX-protective Na_V_1.4 mutations is otherwise paradoxical.

### Evolutionary logic of toxin sequestration over Na_V_ mutation

From an evolutionary perspective, sequestration is an elegant solution. Na_V_s are among the most conserved and key proteins in vertebrate excitable tissues^51,52^. Even small perturbations in Na_V_1.4 gating can impact muscle performance, a trait under strong selection, especially for passerine birds^53^. A lineage that acquires a dietary toxin such as BTX from Melyrid beetles^54^ faces a dilemma: how to keep the toxin for defense but avoid self-harm. Mutating Na_V_1.4 solves only the second part of that problem, and to date, no BTX-bearing animal has been found to perfectly solve this auto-intoxication issue by target mutations. Moreover, similar to humans, birds have multiple Na_V_ isoforms, consistent with the fact that birds, along with mammals, inherited multiple sodium channel gene ancestors from early tetrapods^55,56^. A target mutation strategy would require mutating all Na_V_ isoforms to make them BTX resistant, a process that would most likely add a functional cost to these vital channels, given the delicate location of the BTX binding sites in the channel pore^12^. By contrast, a “toxin sequestration” strategy can preserve Na_V_ function and thus neuromuscular performance and many other Na_V_-related physiological functions. This strategy can also be evolved by duplicating or repurposing existing binding/transport proteins, allowing a low barrier to entry. This “toxin sequestration” strategy can be expressed in tissue- or compartment-specific patterns in key defensive organs such as the skin and the feathers, allowing defense without systemic poisoning. Such an elegant strategy would also explain why strong BTX defenses can co-occur with unmodified, fully functional Na_V_1.4, as we show here.

A limitation is that we tested only Na_V_1.4, the main skeletal muscle Na_V_ isoform implicated in the prior positive selection analysis and a likely target of BTX in muscle^16^. However, in our prior work, we also showed that both the skeletal muscle isoform Na_V_1.4 and the cardiac isoform Na_V_1.5 from the same BTX-bearing bird (*Pitohui uropygialis meridionalis*) is fully BTX-sensitive^26^, supporting the broader conclusion that BTX tolerance in these birds does not generally arise from reduced BTX sensitivity of major excitable-tissue Na_V_s. Nevertheless, we cannot fully exclude the possibility that BTX tolerance could involve target resistance in other Na_V_ isoforms expressed in the nervous system.

Building on this evolutionary context, it is also notable that BTX-bearing passerines are distributed across multiple New Guinean lineages rather than forming a single, tightly clustered “toxic bird” clade. Identified BTX-positive taxa include *Pitohui spp.* (Oriolidae) and *Ifrita kowaldi* (Ifritidae)^14,15^, and recent work extends BTX carriage to additional species placed in other passerine families, including Pachycephalidae and Oreoicidae^16^. This taxonomic distribution is most consistent with multiple acquisitions of BTX carriage and/or toxin-handling traits rather than requiring a single origin of a Na_V_ target-mutation solution. While resolving the number and timing of origins of BTX tolerance will require broader comparative sampling, our data directly test the specific mechanistic claim that the two Na_V_1.4 substitutions identified under positive selection confer autoresistance via target modification and do not support that model^16^. Notably, Bodawatta et al.’s own cross-species Na_V_1.4 alignment shows that D1050N and S1568P also occur in multiple bird species considered non-toxic or not yet tested for BTX^16^, consistent with our functional data that these substitutions alone are insufficient to confer BTX resistance. Together with prior functional evidence that BTX-bearing birds can retain BTX-sensitive Na_V_s^26^, this phylogenetic distribution further reinforces the plausibility of upstream protection mechanisms as a general evolutionary solution. Importantly, the fact that toxin-based chemical defense systems have evolved independently four times in neotropical Dendrobatidae poison frogs, including BTX-carrying *Phyllobates* frogs^13^, and in multiple lineages of toxic birds, including BTX-carrying *Pitohui* and *Ifrita* birds^14,15^, supports the idea that such general “toxin sequestration” mechanisms may underlie toxin autoresistance. Future studies could focus on identifying the specific “protein sponge” used by these BTX-carrying birds.

Finally, our data rule out a Na_V_1.4 target-mutation mechanism for the specific substitutions tested here, but they do not identify the upstream factors that prevent BTX from engaging Na_V_s in vivo. Thus, while our results are consistent with toxin sequestration or compartmentalization models proposed in other BTX-bearing vertebrates, they should not be interpreted as direct evidence for a BTX-binding “toxin sponge.” At present, no BTX-binding protein has been identified in birds, and defining the responsible protective mechanisms will require additional biochemical and physiological work. Plausible candidates include soluble BTX-binding proteins or carrier complexes in circulation, transport pathways that concentrate BTX in skin/feathers, or intracellular compartmentalization that limits free BTX in excitable tissues. Establishing the molecular identity, affinity, and tissue distribution of such factors is an important next step.

### Implications for antidote discovery

If BTX sequestration is indeed the primary strategy in toxic birds and frogs, isolating the responsible binding proteins becomes an obvious translational goal. BTX-binding proteins could be 1) engineered for higher affinity/specificity; 2) expressed recombinantly and used as circulating decoys; or 3) coupled to delivery platforms to clear BTX after exposure. This mirrors efforts in the saxitoxin field, where naturally occurring binding proteins have inspired therapeutic tools^57^. Our work narrows the search space for BTX countermeasure. Because Na_V_1.4 is BTX sensitive in BTX-carrying species, soluble or membranous protective factors must be upstream to prevent the toxin from reaching Na_V_s in the first place.

### Conclusions and Perspectives

Our results refute the notion that Na_V_1.4 mutations confer BTX autoresistance in toxic birds^16^. Instead, these findings strengthen the emerging view that toxin sequestration underlies BTX tolerance across birds and amphibians^26,33^. Future studies should aim to identify the molecular identity and binding properties of BTX-binding proteins, elucidate their expression patterns across tissues, and explore whether similar systems protect other toxin-bearing vertebrates, consistent with the growing list of “toxin-sponge” proteins^28–32^. Understanding this natural defense mechanism could inform the design of synthetic BTX antidotes and molecular sensors and contribute broadly to strategies for neutralizing Na_V_-targeting toxins in clinical and environmental contexts.

## Acknowledgements

We thank M. DiFranco for gifting us batrachotoxin, D.B. Leitch and K.A. Jønsson for comments on the manuscript, and members of the Abderemane Lab at UCLA for insightful discussion. This work was supported by the Cannella & Bianco Charitable Fund to F.A.-A.

## Author Contributions

E.G., Z.J. and F.A.-A. conceived the study and designed the experiments. E.G. performed molecular biology experiments, electrophysiology experiments, and analyzed the data, with help from M.J.T., H.Y., and D.Y. for electrophysiology data acquisition and processing. Z.J. performed structural modeling and analysis. F.A.-A. analyzed data and provided guidance and support. E.G., Z.J. and F.A.-A. wrote the paper.

## STAR METHODS

### KEY RESOURCES TABLE

| REAGENT or RESOURCE | SOURCE | IDENTIFIER |
| --- | --- | --- |
| **Bacterial and Virus Strains** |  |  |
| Dh5a competent E. coli | MCLAB | Cat#DA-196 |
| **Chemicals, Peptides, and Recombinant Proteins** |  |  |
| Potassium chloride | Fisher Chemical | Cat#P217-500 |
| Sodium chloride | Fisher Bioreagents | Cat#BP358-1 |
| Calcium chloride | Sigma Aldrich | Cat#C3306-100G |
| Magnesium chloride | Sigma Aldrich | Cat#208337-100G |
| HEPES (4-(2-hydroxyethyl)-1-piperazineethanesulfonic acid) | Thermo Fisher Scientific | Cat#A14777.30 |
| Sodium hydroxide | Chem Cruz | Cat#SC-203387 |
| penicillin/streptomycin | Gibco | Cat#15140-122 |
| gentamicin | Fisher Bioreagents | Cat#BP918-1 |
| Batrachotoxin | Chem Cruz | Cat#SC-201086 |
| **Critical Commercial Assays** |  |  |
| T7 mMessenger kit | Thermo Fisher Scientific | Cat# AM1344 |
| QuikChange Site-Directed Mutagenesis Kit | Agilent | Cat#200515 |
| **Experimental Models** |  |  |
| Xenopus oocytes | This study | N/A |
| **Recombinant DNA** |  |  |
| Human Nav1.4 | Abderemane-ali et al., 2021 | N/A |
| Pitohui Nav1.4 | Abderemane-ali et al., 2021 | N/A |
| Pitohui Nav1.4 D1049N | This study | N/A |
| Pitohui Nav1.4 S1581P | This study | N/A |
| Pitohui Nav1.4 D1049N/S1581P | This study | N/A |
| Human Navβ1 | Abderemane-ali et al., 2021 | N/A |
| **Software and Algorithms** |  |  |
| pCLAMP 9 | Molecular Devices | N/A |
| Clampfit 10.7 | Molecular Devices | N/A |
| SigmaPlot 12.5 | Grafiti LLC | N/A |
| GraphPad 10 | Prism | N/A |
| ChimeraX 1.9 | UCSF | N/A |
| Jalview 2.11.5.1 | N/A | N/A |

### CONTACT FOR REAGENT AND RESOURCE SHARING

Further information and requests for resources and reagents should be directed to and will be fulfilled by the Lead Contact, Fayal Abderemane-Ali.

### EXPERIMENTAL MODEL AND SUBJECT DETAILS

Oocytes collection: Oocytes were harvested from female *Xenopus laevis* frogs purchased from Xenopus and housed in the UCLA Division of Laboratory Animal Medicine (DLAM) facilities. The use of these Xenopus oocytes was approved by IACUC (protocol approval # ARC-2023-051), and experiments were performed in accordance with University of California guidelines and regulations.

### METHOD DETAILS

#### Protein structure prediction

The full-length structure of *Pitohui uropygialis meridionalis* Na_V_1.4 (*Pum* Na_V_1.4, GenBank accession no. MZ545383.1) was predicted using the online AlphaFold server (https://alphafoldserver.com/) powered by AlphaFold3^58^. Model confidence was evaluated by the pLDDT value, which shows a per-atom confidence metric based on the expected local structural accuracy. Regions with pLDDT ≥ 70 were considered reliable.

#### Structure Analysis

Molecular structures were visualized and analysed using UCSF ChimeraX (version 1.10.1)^59^. The AlphaFold-predicted structure of *Pum* Na_V_1.4 was visualized in UCSF ChimeraX. The residues were colored according to their pLDDT values using the “render/select attribute” function, based on four discrete categories: pLDDT<50 (orange), 50-70 (yellow), 70-90 (cyan), and >90 (blue).

The predicted structure of *Pum* Na_V_1.4 was compared with the published structure of human Na_V_1.4 (PDB 6AGF), rat Na_V_1.5 in complex with BTX (PDB 8T6L), human Na_V_1.7 in complex with Na_V_β1 and Na_V_β2 (PDB 7W9L), and human Na_V_1.1 in complex with Na_V_β4 (PDB 7DTD). Protein structures were superimposed using the “Matchmaker” tool, which aligns two structures based on pairwise sequence alignment followed by least-squares fitting of Cα atoms. The published structure was used as a reference to quantify structural similarities. Root-mean-square deviation (RMSD) values were calculated, and the *Pum* Na_V_1.4 structure was colored according to the RMSD values, with a blue-white-red gradient representing RMSD values from 1 to 3 Å. Regions of the channel that are missing in experimental data are colored in gray. The distances between two mutation positions in *Pum* Na_V_1.4 (D1049, S1581) and the reported two BTX binding sites in rat Na_V_1.5 were measured by the “Distance” tool.

#### Site-directed mutagenesis

For electrophysiology experiments, *Hs* Na_V_1.4 (GenBank accession no. NM_000334.4), human Na_V_β1 (GenBank accession no. NM_001037.5), and *Pum* Na_V_1.4 (GenBank accession no. MZ545383.1), all in pCDNA3.1, were used. All mutants were made using Site-Directed Mutagenesis. The mutations were validated by full-plasmid sequencing, including the genes encoding for the proteins of interest.

#### Two-electrode voltage clamp electrophysiology

Two-electrode voltage-clamp recordings were performed on defolliculated stages V–VI *Xenopus laevis* oocytes harvested (under UCLA IACUC protocol ARC-2023-051) and microinjected with mRNA. Linearized cDNAs were translated into capped mRNA using the mMESSAGE mMACHINE T7 and SP6 Transcription Kits (Invitrogen). Oocytes were injected with 0.3 to 0.9 ng of mRNA with a 4:1 alpha:beta1 weight ratio for all conditions. Two-electrode voltage-clamp experiments were performed 2-4 days after injection. Data were acquired using a GeneClamp 500B amplifier (Axon Instruments) controlled by pClamp software (Molecular Devices) and digitized at 1 kHz using a Digidata 1332A digitizer (Axon Instruments). Oocytes were impaled with borosilicate recording microelectrodes (0.3–3.0-MΩ resistance) backfilled with 3 M KCl. Sodium currents were recorded using a bath solution containing the following in mM: 96 NaCl, 1 CaCl2, 1 MgCl2, 2 KCl, and 5 HEPES (pH 7.5 with NaOH) supplemented with antibiotics (50 µg ml−1 gentamicin, 100 IU ml−1 penicillin, and 100 µg ml−1 streptomycin). BTX was applied from the intracellular side of the membrane by injecting oocytes with 50 nL BTX (Santa Cruz Biotechnology, Inc.) at a stock concentration of 1mM for a target intracellular concentration of 50 μM. After BTX injection, oocytes were stimulated by applying 1,000 step pulses of 60 ms each, from −120 to 0 mV at 2 Hz frequency, to facilitate BTX access into the channel pore.

To generate the activation curve, from a holding potential of −120 mV, the membrane was depolarized to values between −100 mV and +50 mV (+5 mV increment) for 60 ms, every 5 s. Sodium current was calculated after leak subtraction. To generate the inactivation curve, from a holding potential of −120 mV, the membrane was depolarized to values between −100 mV and 0 mv (+5 mV increment) for 600 ms, followed by a 60-ms test pulse to 0 mV, every 10 s. Activation and inactivation curves were fitted by the Boltzmann equations. G/V curves are obtained as follows: GNa was calculated from GNa = INa/(V − Vrev), where INa is the peak sodium current, V is the membrane potential and Vrev is the reversal potential estimated for each cell by linear regression of the linear rectification of I/V curve, when channels are fully activated. GV curves were subsequently obtained by dividing at each potential the peak current by the corresponding value of the linear regression curve. To assess recovery from fast inactivation, from a holding potential of −110 mV, the membrane was depolarized with a 2-step depolarization at 0 mV for 60-ms each (control pulse and test pulse), with various times (Δt) between these pulses. Δt = 20, 25, 30, 40, 80, 160, 320, and 640 ms. Fast inactivation kinetics and recovery from fast inactivation were fitted with a mono-exponential decay.

#### Data analysis

Data is analyzed by GraphPad 10 (Prism), Clamfit 10.7 (Molecular Devices), Excel (Microsoft), SigmaPlot 12.5 (Grafiti LLC), ChimeraX 1.9 (UCSF), and Jalview 2.11.5.1. In all experiments, n represents the number of different *Xenopus oocytes* tested. All data are presented as mean ± standard error of the mean. Statistical differences between samples were determined using Student’s t-tests when data were normally distributed and rank-sum tests (Mann-Whitney test) when data were not normally distributed. P-values are calculated relative to *Pum* NaV1.4 or *Pum* NaV1.4 + BTX. A value of P < 0.05 was considered significant.

## Supplementary material

Supplementary material contains Figures S1, S2, and S3.

