## Supplemental File for "Batrachotoxin Sensitive Sodium Channels in Toxic Birds Challenge “Target Mutation” Strategy of Toxin Autoresistance"

5 May 2026

### Figure S1

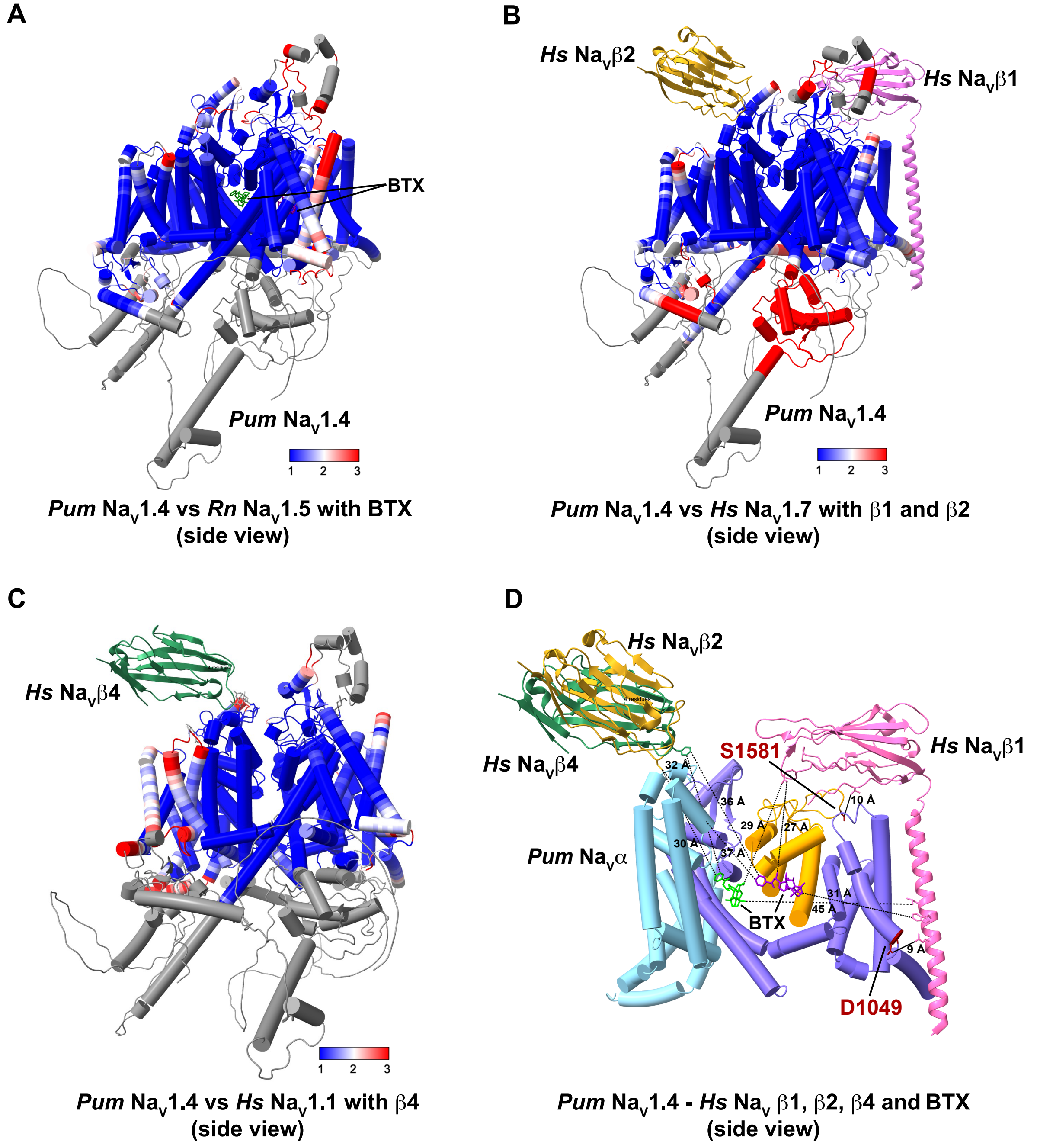
A
BTX
Pum Nav1.4
1
2
3
Pum Nav1.4 vs Rn Nav1.5 with BTX
(side view)
B
Hs Navβ2
Hs Navβ1
Pum Nav1.4
1
2
3
Pum Nav1.4 vs Hs Nav1.7 with β1 and β2
(side view)
C
Hs Navβ4
1
2
3
Pum Nav1.4 vs Hs Nav1.1 with β4
(side view)
D
Hs Navβ2
Hs Navβ4
S1581
Hs Navβ1
32 Å
36 Å
10 Å
29 Å
27 Å
30 Å
37 Å
Pum Navα
31 Å
45 Å
BTX
9 Å
D1049
Pum Nav1.4 - Hs Nav β1, β2, β4 and BTX
(side view)

**Fig. S1: Structural comparison between *Pum* $Na_V$1.4 and other BTX- or $Na_V\beta$-containing $Na_V$ structures. A,** Structural comparison between *Pum* $Na_V$1.4 and *Rn* $Na_V$1.5-BTX. Using *Rn* $Na_V$1.5-BTX (PDB 8T6L) as reference, *Pum* $Na_V$1.4 is colored by per-residue RMSD with a scale ranging from 1 Å (blue) to 3 Å (red). Regions missing in the experimental *Rn* $Na_V$1.5 structure but present in the predicted *Pum* $Na_V$1.4 structure are in light gray. Two BTX molecules are colored in lime and purple, respectively. **B,** Structural comparison between *Pum* $Na_V$1.4 and *Hs* $Na_V$1.7-$Na_V\beta$1-$Na_V\beta$2. Using *Hs* $Na_V$1.7-$Na_V\beta$1-$Na_V\beta$2 (PDB 7W9L) as reference, *Pum* $Na_V$1.4 is colored by per-residue RMSD with a scale ranging from 1 Å (blue) to 3 Å (red). Regions missing in the experimental *Hs* $Na_V$1.7-$Na_V\beta$1-$Na_V\beta$2 structure but present in the predicted *Pum* $Na_V$1.4 structure are in light gray. *Hs* $Na_V\beta$1 and $Na_V\beta$2 are colored in pink and gold, respectively. **C,** Structural comparison between *Pum* $Na_V$1.4 and *Hs* $Na_V$1.1-$Na_V\beta$4. Using *Hs* $Na_V$1.1-$Na_V\beta$4 (PDB 7DTD) as reference, *Pum* $Na_V$1.4 is colored by per-residue RMSD with a scale ranging from 1 Å (blue) to 3 Å (red). Regions missing in the experimental *Hs* $Na_V$1.1-$Na_V\beta$4 structure but present in the predicted *Pum* $Na_V$1.4 structure are in light gray. *Hs* $Na_V\beta$4 is colored in green. **D,** Structural model of *Pum* $Na_V$1.4-BTX-*Hs* $Na_V\beta$1/$\beta$2/$\beta$4. *Hs* $Na_V\beta$1, $Na_V\beta$2 and $Na_V\beta$4 are colored in pink, gold, and green, respectively. Residues D1049 and S1581 are colored in red and indicated. BTX molecules inside the channel inner cavity are colored in lime and purple. Distances between BTX molecules and $Na_V\beta$ subunits are indicated in dashed lines. For clarity, only parts of Domains II, III, and IV are shown, and Domain I is omitted.

### Figure S2

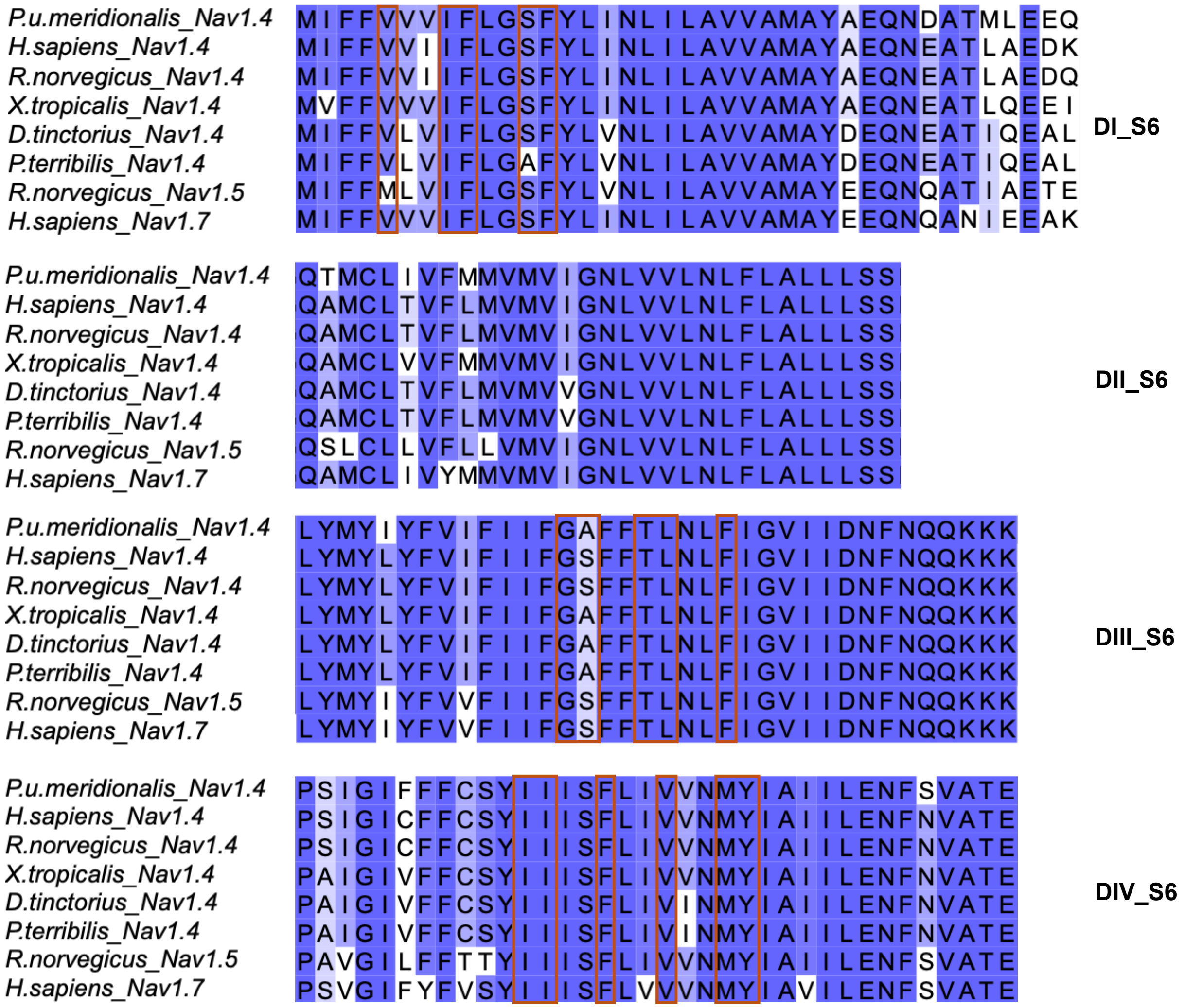


**Figure S2 Conservation of pore lining segments and BTX-interacting residues among $Na_V$ orthologs used for structural mapping or from toxic and non-toxic species.** S6 Sequence alignment of *Pitohui uropygialis meridionalis* $Na_V$1.4 (QYB47537.1), *Homo sapiens* $Na_V$1.4 (NP_000325.4), *Rattus norgevicus* $Na_V$1.4 (NP_037310.1), *Xenopus tropicalis* $Na_V$1.4 (XP_017944811.2), *Dendrobates tinctorius* $Na_V$1.4 (QYB47541.1), *Phyllobates terribilis* $Na_V$1.4 (QYB47540.1), *Rattus norgevicus* $Na_V$1.5 (NM_013125.4), and *Homo sapiens* $Na_V$1.7 (Q15858.3). Residues conserved in all constructs are colored in blue. Residues interacting with BTX in the *Rn* $Na_V$1.5-BTX structure (PDB 8T6L) are highlighted (red boxes). S6 segments are labeled using boundaries from (36).

### Figure S3

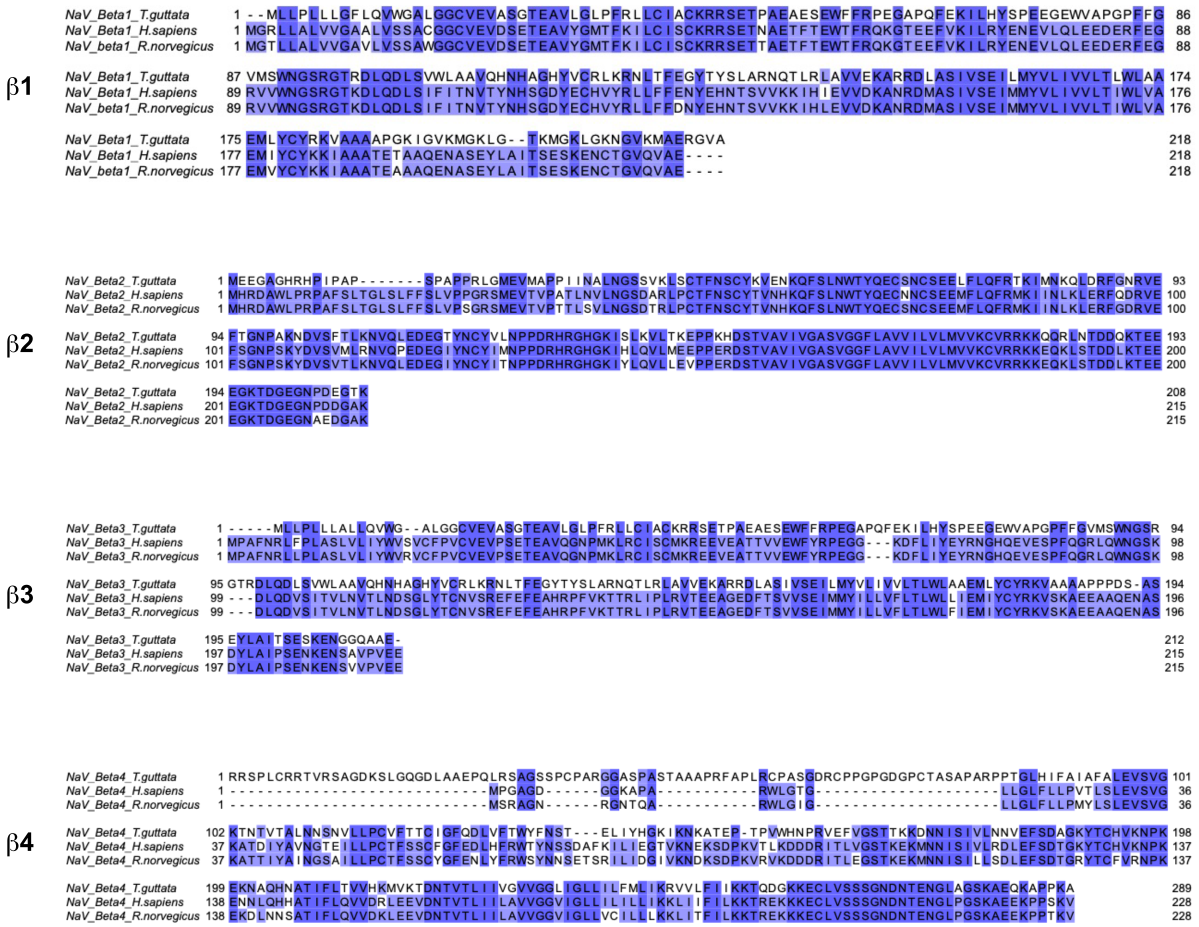


**Figure S3 Conservation of Na$_V$β subunits (β1, β2, β3, and β4) among *Taeniopygia guttata*, *Homo sapiens, and Rattus norgevicus*.** Sequence alignment of *Tg* Na$_V$β1 (A0A674GBS6), *Tg* Na$_V$β2 (H1A422), *Tg* Na$_V$β3 (A0A674HLS7), and *Tg* Na$_V$β4 (H0YPN3); *Hs* Na$_V$β1 (Q07699), *Hs* Na$_V$β2 (O60939), *Hs* Na$_V$β3 (Q9NY72), *Hs* Na$_V$β4 (Q8IWT1); and *Rn* Na$_V$β1 (Q00954), *Rn* Na$_V$β2 (P54900), *Rn* Na$_V$β3 (Q9JK00), and *Rn* Na$_V$β4 (Q7M730). Residues conserved in all constructs are colored in blue.
